# Unified Analysis of Multi-order Tensors for Integrative Molecular Profiling

**DOI:** 10.1101/2024.11.04.621951

**Authors:** Kevin De Azevedo, Florian Buettner

**Affiliations:** Goethe University, Frankfurt, Germany; German Cancer Research Center (DKFZ), Germany; German Cancer Consortium (DKTK), Frankfurt, Germany; Frankfurt Cancer Institute, Germany

**Keywords:** Tensor decomposition, Bayesian Inference, Factor analysis

## Abstract

In recent years, the exponential growth of high-dimensional, multi-modal molecular data has created both opportunities and challenges in personalized medicine. While existing approaches like matrix decomposition and neural network-based embeddings have been used to analyze such data, they have limitations in interpretability, handling missing values, and treating features across modalities as unrelated. To address these challenges, we present MUSIC (MUltiview BayeSIan Tensor DeComposition), a novel framework for probabilistic multi-view tensor decomposition that can integrate collections of tensors of different orders. MUSIC combines the strengths of group factor analysis and tensor decomposition through a Bayesian approach with structured sparsity priors. The framework offers several key advantages: (1) fast model training using variational inference, (2) inference of interpretable embeddings via structured sparsity, (3) efficient handling of missing values, and (4) flexible combination of tensors of different orders. We demonstrate MUSIC’s effectiveness on both simulated data and real-world applications, including drug response analysis in CLL patients and multi-modal single-cell data analysis in leukemia patients. Our results show that MUSIC can reveal interpretable multi-modal patterns capturing structured variation across patients, cell types, and modalities that are associated with disease states and can be explained through cell type- and modality-specific pathway activities.

## 1 Introduction

In recent years, the exponential growth of high-dimensional, multi-modal molecular data has presented both opportunities and challenges across applications in personalised medicine, where the comprehensive characterisations of patients across molecular layers has revealed resistance mechanisms and prognostic subgroups across entities [9, 12, 14, 18]. These multi-modal approaches are most promising in settings where outcome heterogeneity between individuals is largely unexplained. With the primary sources of inter-patient differences being unknown, unsupervised modelling approaches are needed that can reveal the main axes of variation in an interpretable manner. Common approaches typically build on matrix decomposition via principal component analysis or group factor analysis (e.g. as implemented in the commonly used MOFA model) [1]. Alternatively, neural network based embeddings have been proposed [7]. However, the latter has the drawback of inferring embeddings with limited interpretability so that it is often not clear what biological processes are being captured. In addition, neural network based approaches are often not able to account for missing values - a common occurrence in multi-omics studies, where not all modalities can be profiled for all patients - in a principled manner [1, 17]. Both paradigms suffer from an inherent drawback in that they treat features in each modality as unrelated. In many setups however, there is a direct one-to-one correspondence between features of different data views, for example when different tissues of a patient are being profiled via single-cell experiments [16] or in the case of drug-response experiments, where in a cohort of patients response to drugs is being quantified in different conditions [3]. Most, recently, Mitchell et al [16] have proposed to generalise PCA-style matrix decomposition to third order tensors and introduced a tensor decomposition approach to jointly model expression variation across multiple cell types. However, this approach is limited to data that can be represented in a single tensor (in this case: patients x genes x tissues). In many applications, however, a natural representation of multimodal data is via a collection (list) of tensors of different order. This can for example be a drug response tensor of third order (patients x drugs x conditions) in addition to RNA-seq data in from of a second-order tensor (patients x genes). Similarly, in case of modeling inter-patient variation across tissues, patients are often not only profiled in terms of gene expression as described above (patients x genes x tissues), but also via additional modalities such as ATAC-seq to measure chromatin accessibility [5].

Here, we propose MUSIC, a framework for probabilistic multi-view tensor decomposition that can integrate a collection of tensors of different order. This can be interpreted as a generalisation of group factor analysis to tensors of varying order or, orthogonally, as a generalisation of tensor decomposition approaches such as PARAllel FACtor analysis (PARAFAC) to multiple views. More specifically, we propose a Bayesian approach to integrate multiple tensors via structured sparsity priors based on view-wise automatic relevance determination (ARD) [13]. We demonstrate on simulated data that our model is able to effectively model collections of tensors of different orders and with different ratios of missing values and falls back to tensor composition with the same efficacy as standard PARAFAC in the special case of only one tensor. We then demonstrate on different real-world datasets the diverse applicability of MUSIC. First, we use it to disentangle the variability of drug-response data of 192 CLL patients who were profiled with RNA-seq (first data view, tensor 2nd order) and for whom the influence of 17 cytokines on 12 drugs was quantified (second data view, tensor 3rd order) [3]. We show that MUSIC is able to learn interpretable representations that (i) recover the correlation structure between drugs and (ii) that reveal patient subgroups associated to clinically relevant covariates. In a second usecase, we employ MUSIC to identify multi-cellular and multi-modal patterns characterizing patients with KMT2A-rearranged leukemia and healthy donors who were profiled with single-cell RNA-seq and single-cell ATAC-seq. A natural representation of these data is in form of two tensors of third order, with the first tensor capturing variability across patients, genes and cell types and the second tensor capturing variability across patients, peaks and cell types. This revealed interpretable multi-modal patterns capturing structured variation across patients, cell types and modalities that were associated to disease state and could be explained via cell type- and modality-specific pathway activities.

## 2 Material and Methods

### 2.1 Model formulation

At its core, MUSIC builds upon the statistical frameworks of group factor analysis on the one hand and tensor decomposition on the other hand. Drawing from both modeling approaches, we introduce a Bayesian group tensor decomposition framework that is tailored to the requirements of multi-modal studies: (i) fast model training using variational inference, (ii) inference of interpretable embeddings via structured sparsity, (iii) efficient handling of missing values and (iv) flexible combination of tensors of different orders, allowing for the joint modeling of diverse combinations of modalities. Special cases of MUSIC are group factor analysis as implemented in MOFA [1] (combination of rank-2 tensors only) and tensor decomposition such as PARAFAC or the recently proposed scITD model [16] (one rank-3 tensor only).

#### Model Specification

In the following, we specify the generative model. The full generative model including hyperparameter choices along with the joint probability density function is detailed in the appendix. Let *G* = {**Y**^**1**^, … **Y**^**M**^} be a collection of tensors of rank 2 and rank 3 respectively. Let 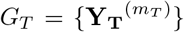 be the subset of *M*_*T*_ three-way tensors of dimension 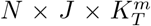, with *N* samples (e.g. patients), *J* slices (e.g. tissues) and 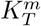 features (e.g. genes); let *G*_*M*_ = {**Y**_**M**_^(*m*)^} be the subset of two-way tensors of dimension 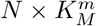 of *N* samples (e.g. patients), and 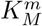 features (e.g. genes) such that *G*_*T*_ ∪ *G*_*M*_ = *G*. Note that all tensors in *G*_*T*_ share the number of samples *N* and number of slices *J*, but will have different sets of features (e.g. gene expression in one tensor and peaks from ATAC-seq in a second tensor); all two-way tensors (matrices) in *G*_*M*_ share the same number of samples *N* across all tensors in *G*, but have different features. MUSIC then jointly decomposes the tensors as:

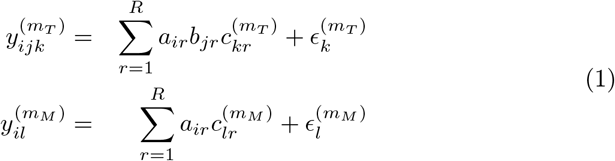

where **a**_*i*_ is the *R*-dimensional embedding of the *i*th sample^5^ (shared across all tensors in *G*), **b**_*j*_ is the *R*-dimensional embedding of the *j*th slice (shared across all tensors in *G*_*T*_) and 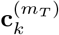 and 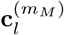 correspond to the view-specific *R*-dimensional embeddings of features for tensors in *G*_*T*_ and *G*_*M*_ respectively. 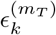 and 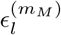 denote the feature- and view-specific residual noise terms. We assume a Gaussian noise model with variance 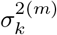 and place a conjugate inverse Gamma prior on the variance and normal priors on the sample- and slice-embeddings (see Supplement for details).

#### Structured Sparsity

To disentangle view- and sample-specific heterogeneity, we use a multi-scale shrinkage prior on **C**^(*m*)^ (the view-specific feature embedding in matrix form) to enable global and local sparsity. We encourage this structured sparsity via a Horseshoe Prior that is amenable to variational inference [4, 17]. More specifically, we model **C**^(*m*)^ as follows:

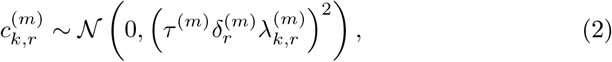

where where we place a positive Cauchy prior on each scale,

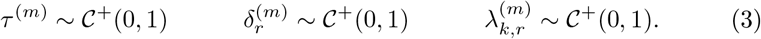

Here, 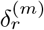 can be interpreted as an automatic relevance determination (ARD) prior [13] for the *r*th factor in view *m*, which decouples this factor from the other views, thereby allowing for individual factors to only explain variance (i.e. be active) in a subset of views (or tensors). On an element-wise level, 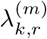 regularizes each individual loading, encouraging element-wise shrinkage and sparsity.

#### Variational Inference

We approximate the posterior distributions of the variables of interest **Θ** by introducing a fully factorized family of parameterized distributions, where **Θ** = **A, B, C, Λ, Δ**, *τ*, **Ψ**, and

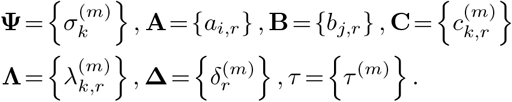

This is achieved by maximizing the evidence lower bound with respect to variational parameters. This, in turn, reduces the gap between the true and the approximate posterior in terms of the KL divergence. We use the family of normal distributions for approximating **C**^(*m*)^, **A** and **B**^(*m*)^; for the rest of the parameters we use the log-Normal distribution to ensure positive samples [8]. We provide an implementation of MUSIC in Pyro [2] that can be accessed at https://github.com/MLO-lab/music/tree/dev.

### 2.2 Datasets

We demonstrate the versatility of MUSIC to disentangle inter-patient heterogeneity in complex study designs via two publicly available datasets:

– A dataset comprised of RNA-seq and viability assays of cell lines of patients suffering from chronic lymphocytic leukemia (CLL) [3]. We use the log-normalized survival values of the cell lines treated with 17 different drugs and 12 different cytokine stimuli, as well as the RNA-seq data of the CLL patients, taking the 5000 most variable features and log-normalizing the values. We construct a rank-3 tensor of patients x drugs x features and in addition a matrix of patients x genes representing the RNA-seq data.
– A single-cell multi-omics experiment on 18 pediatric patients with acute lymphoblastic leukemia (ALL) supplemented with 5 healthy individuals [5]. We use the single-cell RNA-seq and single-cell ATAC-seq data available, eliminating cells with a total count of RNA below 200, and taking the 5000 most variable features in each case. We use the Seurat-normalized values provided in the dataset.

We also construct synthetic data for further evaluation. Briefly, we create normally-distributed embedding matrices that are multiplied as specified in equation (1) to form a set of 3-dimensional tensors.

### 2.3 Experiments

#### Cytokine experiment

Our goal for this experiment is to evaluate the added value of jointly modeling the drug-response data with the RNA-seq data. We therefore first run a single-view version of MUSIC based only on the drug-response tensor. We then run the multi-view version, jointly decomposing the drug-response tensor and the RNA-seq data matrix. We run both versions with *R* = 20 latent factors and compute factor loadings, by performing batch matrix multiplication of the corresponding factor matrices. This results in three factor loadings respectively of size Patients x Drug X R, Patients x Cytokine X R, and Drug x Cytokine x R.

#### Single cell experiment

In this experiment we aim to demonstrate the ability of MUSIC to disentangle patient- and cell-type specific variation from complex single-cell multi-omics data. Chen et. al [5] data provide the cell types of each cell within the data. We follow [16] and summarize the normalized values across patients and across cell types, to obtain a pseudo-bulk representation of the data. We then obtain two tensors of dimension patients x cell types x features, one for each data modality. We run those with the model with 3000 epochs and a learning rate of 0.005. Next, to interpret the sources of variability captured by individual factors, we multiply the cell types and features factor matrices to obtain cell types loadings relatives to each feature. We apply hierarchical clustering on these generalized loadings and extract the linkage matrix. We analyze these clusters by extracting the top features, and using them in over-representation analysis using WebGestalt [6].

#### Synthetic experiment

To demonstrate that our implementation yields competitive results in terms of data reconstruction, we compare a single-view version of MUSIC to a state-of-the-art implementation of PARAFAC as provided as part of the Tensorly package [10]. We therefore create several synthetic single-view tensor dataset for varying number of samples and additionally varying number of seeds. The total combinations of seeds and samples is given by Table 1. We then decompose these simulated tensors with our model and Tensorly’s PARAFAC method, and compute the RMSE between the inferred tensor and the simulated tensor.

**Table 1:** WebGestalt results for pDC cluster transcripts.

| GO Term | Description | Ratio | P-value | FDR |
| --- | --- | --- | --- | --- |
| GO:0002250 | adaptive immune response | 1.9461 | 3.1826e-7 | 0.00018499 |
| GO:0007159 | leukocyte cell-cell adhesion | 2.0238 | 3.3122e-7 | 0.00018499 |
| GO:0002697 | regulation of immune effector process | 1.9825 | 1.6315e-6 | 0.00060747 |
| GO:0022407 | regulation of cell-cell adhesion | 1.8432 | 2.4894e-6 | 0.00069516 |
| GO:0140375 | immune receptor activity | 2.6115 | 5.5682e-6 | 0.0010960 |
| GO:0002683 | negative regulation of immune system process | 1.7948 | 5.8872e-6 | 0.0010960 |
| GO:0051249 | regulation of lymphocyte activation | 1.7618 | 1.2461e-5 | 0.0019530 |
| GO:0002443 | leukocyte mediated immunity | 1.8672 | 1.3987e-5 | 0.0019530 |
| GO:0023023 | MHC protein complex binding | 4.4501 | 1.8502e-5 | 0.0022964 |
| GO:0050900 | leukocyte migration | 1.8393 | 2.5371e-5 | 0.0028339 |

#### Comparison with MOFA

To demonstrate the added value of jointly modelling cytokines and drugs in tensor-form, we compare MUSIC’s performance against another multi-omics integration tool that is based on group factor analysis, namly MOFA [1]. We run MOFA on the CLL data with the same number of factors used for MUSIC, and using the same normalization steps. Since MOFA is not built to deal with tensors, we instead concatenate the viability data with the transcriptome data. We consider the features to be the drugs and the transcripts, and the views the different type of cytokine treatment with the added RNA view. That means that in contrast to MUSIC each cytokine has to be modelled as independent view.

## 3 Results

### 3.1 Reconstruction error

We find that single-view MUSIC is comparable to a state of the art PARAFAC implementation when it comes to reconstruction error. It can reach similar values of RMSE as Tensorly’s PARAFAC method on the same synthethic dataset for different choices of *R* (Fig. 1). In contrast to PARAFAC, however, the shrinkage priors of MUSIC tend to yield sparser solutions, allowing for better interpretability. In addition, PARAFAC is only able to decompose a single tensor and does not allow for missing values.

**Fig. 1:**
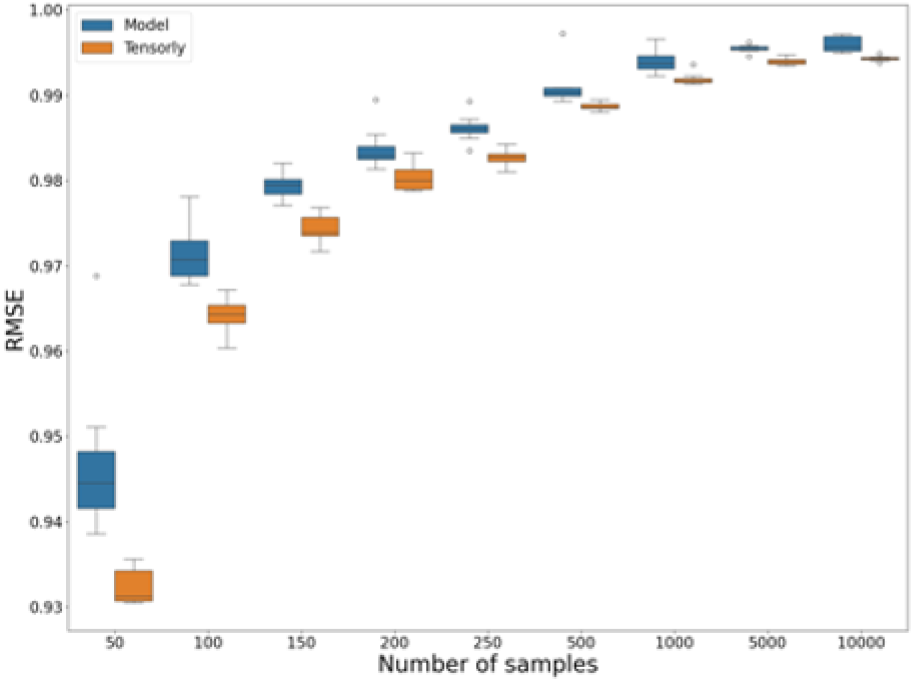
RMSE for varying number of samples for our model and Tensorly’s Parafac method. Note that while especially for smaller sample sizes there is a systematic difference in RMSE, the absolute difference in terms of reconstruction error is very small with values typically below 0.01.

### 3.2 Cytokine data

To inspect the added value of the RNA data to the model, we want to investigate two things: first, if the intrinsic biological correlation structure of the data is maintained, and second if meaningful factors can be inferred by adding the RNA view. For the first point, we look to see if the correlation between the drugs are maintained in the generalized loadings of the patients and drugs when decomposing the drug response tensor alone or when running MUSIC jointly on the drug response and RNA-seq data. In particular, we assess if the drugs involved in the BCR pathways (ibrutinib, idelalisib, PRT062607, IBET-762 and selumetinib) are correlated in the latent space. When looking at the linkage of the hierarchical clustering (Fig. 2) of the drug embeddings, we can observe that the association are similar in both cases, and the drugs cluster closely together with the exception of ibutrinib, showing that the biological association are conserved. When assessing the patient embeddings, only MUSIC run on the multimodal data (drug response and RNA-seq data) jointly is able to infer meaningful embeddings that are associated with two relevant clinical covariates, namely IGHV status and Trisomy 12 status (Fig. 3), better than other methods such as MOFA (Fig. 4). This illustrates that MUSIC is able to leverage data from additional views to infer more meaningful embeddings on the patient level, without compromising on the quality of the drug embeddings.

**Fig. 2:**
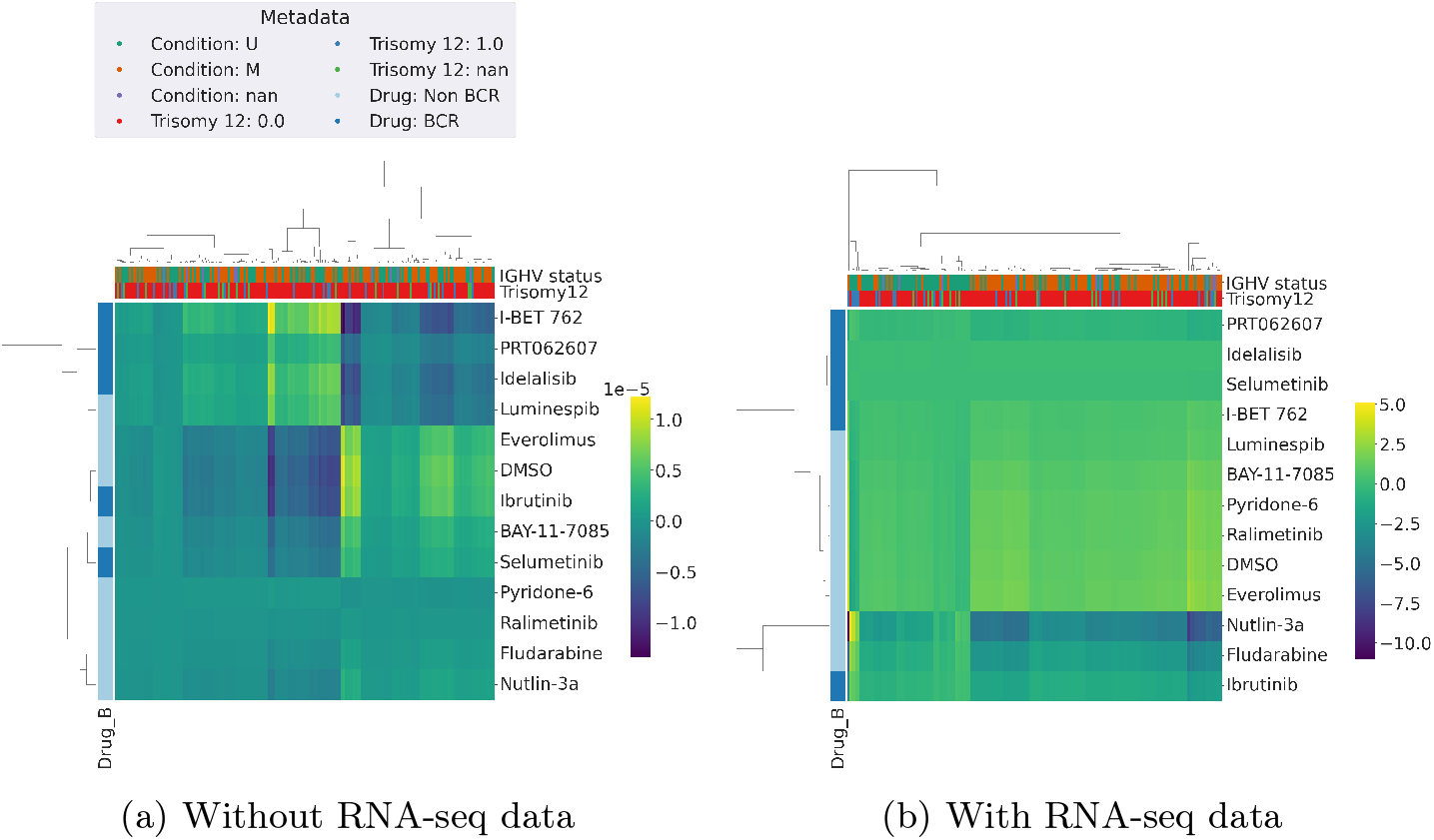
Heatmap of the factor loadings for the patients relative to the drugs for the first factor. The x axis represent the patients

**Fig. 3:**
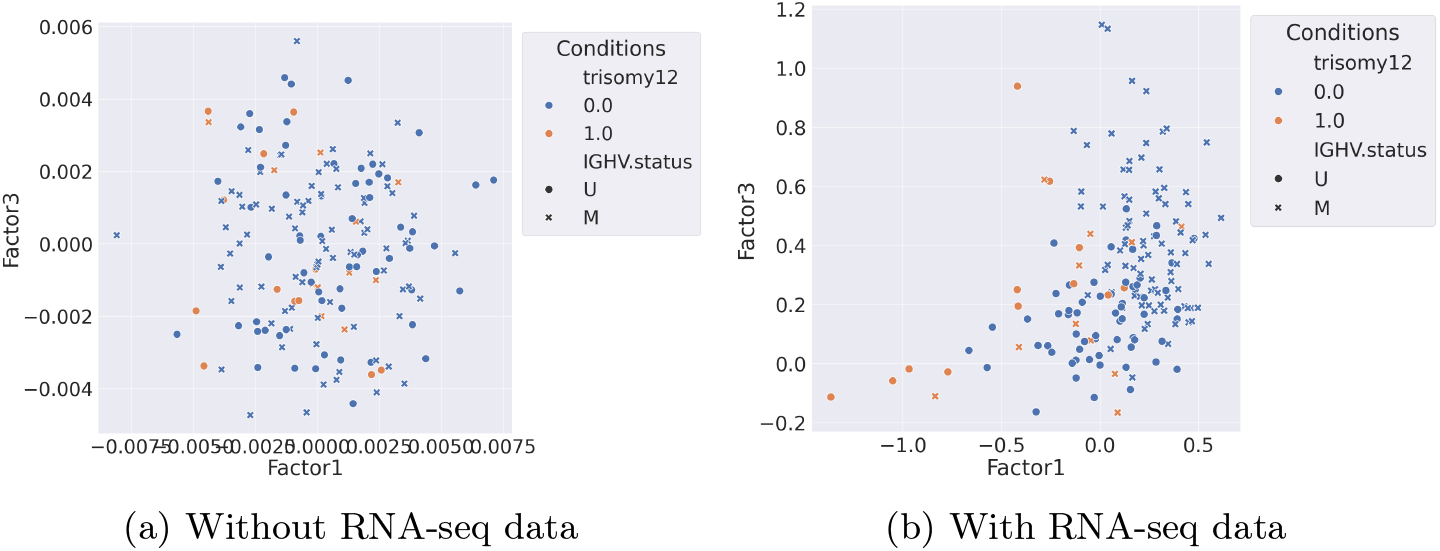
Patient embeddings in the dataset without RNA (left) and with RNA (right). Patients are represented by their IGHV status and their Trisomy 12 status

**Fig. 4:**
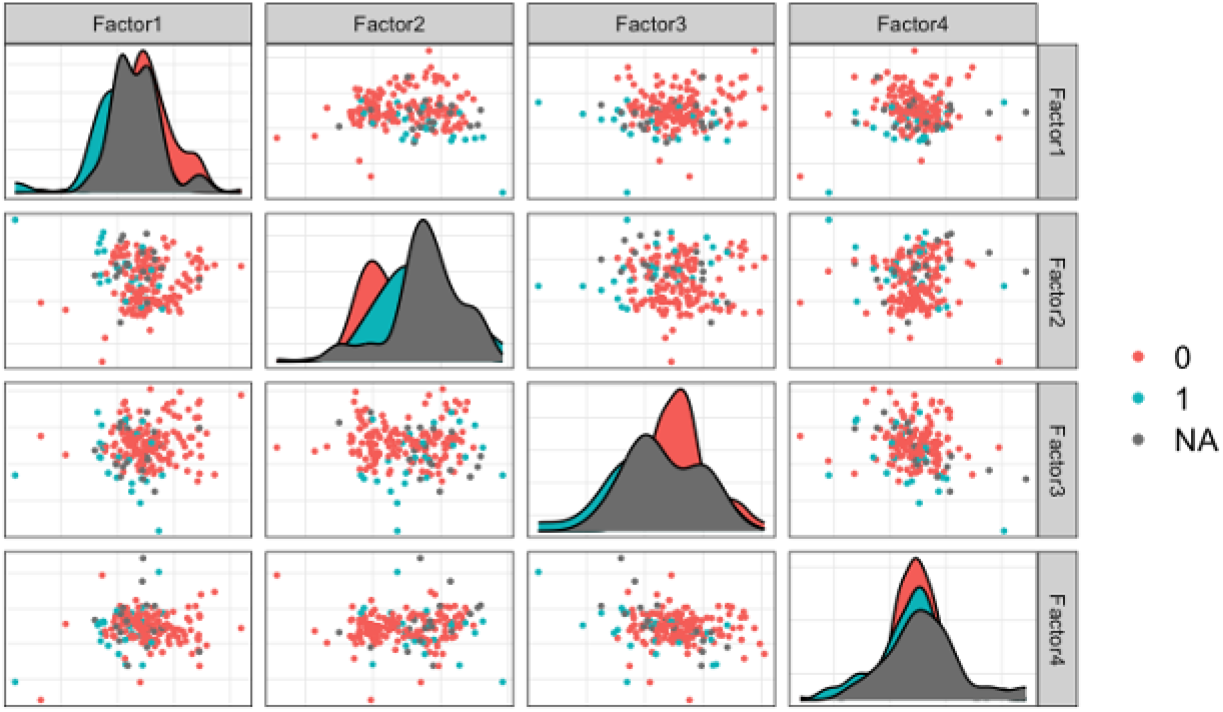
Patient embeddings in the dataset with RNA after being run with MOFA

### 3.3 Single cell data

With the ALL data, we are interested in identifying modules of features in the latent space that jointly characterize variability between patients cell types, both in the scRNA-seq and the scATAC-seq data. Firstly, the model is able to correctly separate healthy and cancer samples in the patients embeddings (Fig. 5). These patient embeddings reveal however that a small subset of ALL patients cluster together, and differ from other patients mostly by their factor values in Factor 5 (Fig. 5). The generalized cell-type/feature loadings matrix associated to this factor reveals that factor 5 is mainly active in pDC cells, both for transcriptomics or epigenetics features (Fig. 6). The result of the over-representation analysis of the genes involved in that separation show pathways associated with pDC cell physiology. For instance, we see pathways associated with MHC activity and adaptive response. pDC are involved in adaptive response by maturing after contact with a virus, and are involved in the cascade of activation of MHC Class 1 and 2 [15]. This illustrates that, MUSIC is able to identify multi-modal feature modules that explain the biological variability between cellular and biological components via it’s structured sparsity prior.

**Fig. 5:**
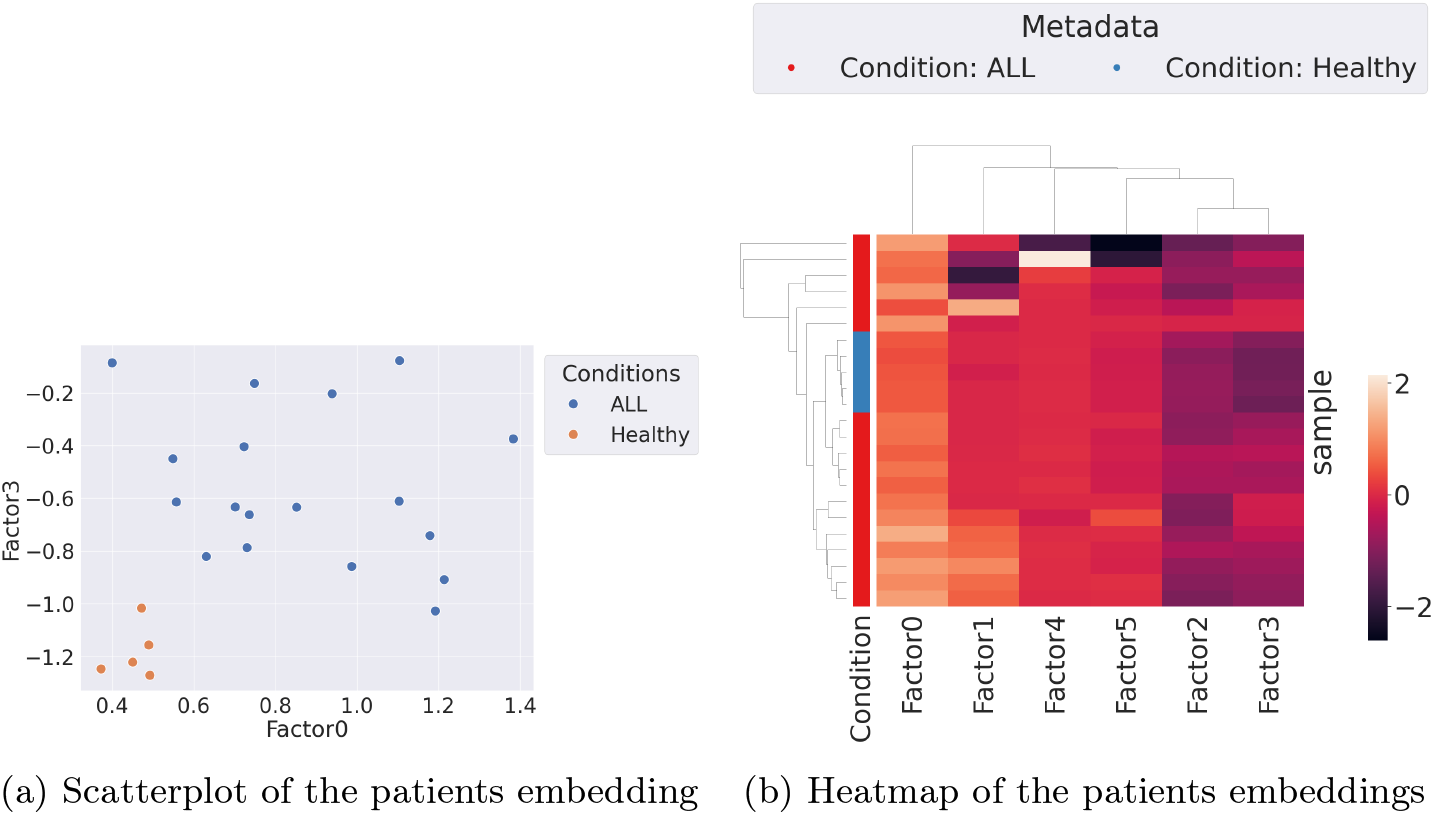
Heatmap and scatterplot of the patient embeddings

**Fig. 6:**
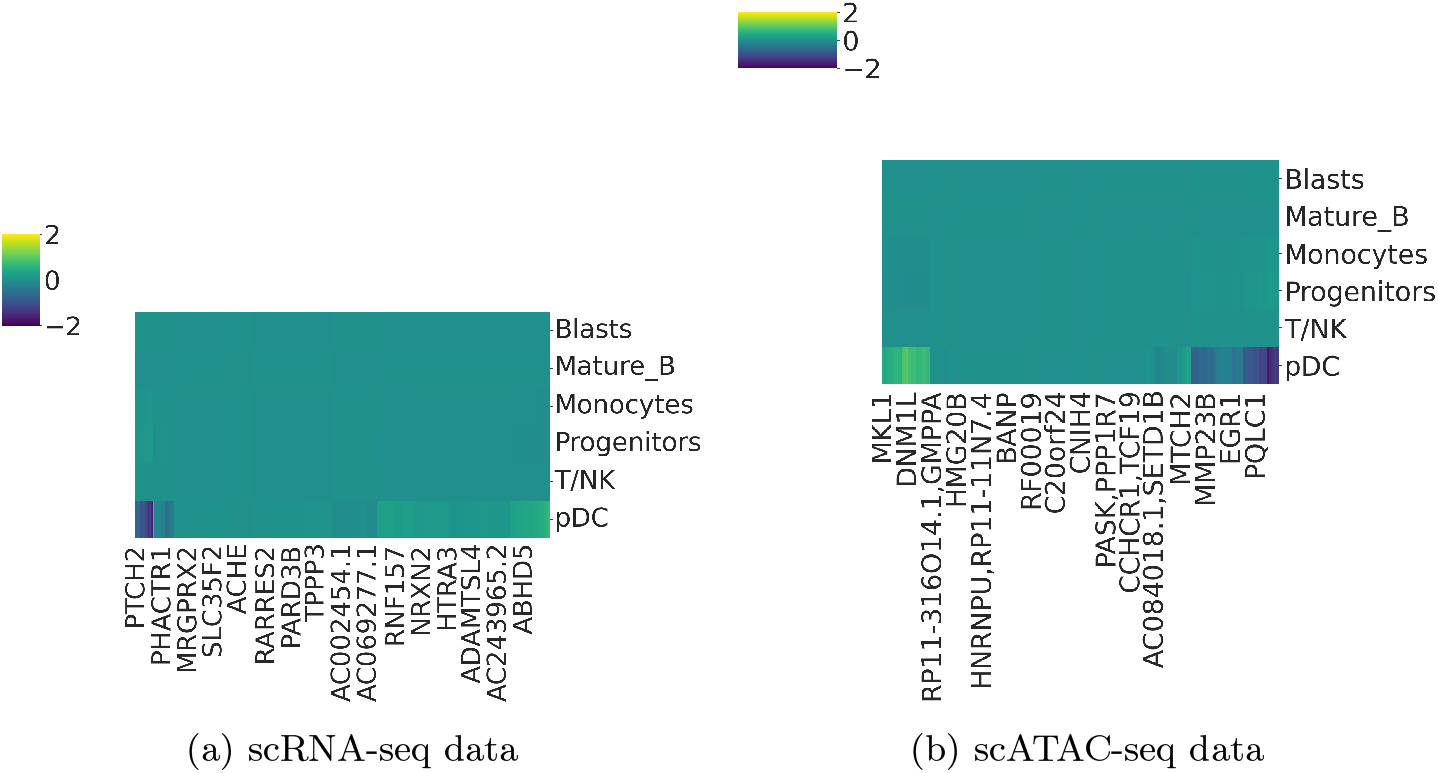
Heatmap of the factor loadings for the cell types relative to the features for factor 5. The x axis represent features.

We show additional generalised loadings plots of Factors 0 and Factor 3 in the Supplement, to further illustrate the broad biological processes driving the separation in terms of healthy donors/cancer patients across multiple cell types, in particular Blasts, Progenitors and T/NK cells.

## 4 Discussion and Conclusion

We have developed MUSIC, a multi-view integration tool that can work with complex study designs to explain the biological variability between patients, with diverse usecases ranging from drug-response studies to single-cell multi-omics data. MUSIC is versatile in that any kind of features can be used in conjunction, without the necessity of having to choose between mapping features to each other as required by tensor decomposition tools and treating all sets of features as independent as in group factor analysis based approaches. We demonstrate the broad applicability of MUSIC on two real-worked datasets and show that the ability jointly decompose tensors of different ranks is crucial to identify patient embeddings that are associated to relevant clinical covariates.

MUSIC is an unsupervised model and as such able to identify multicellular patterns with high variance across patients. However, this means that not all factors will be linked to known covariates and while the extension of this work towards supervised factorizations is in principle possible via the introduction of additional terms to the loss function [11], this comes with new challenges around overfitting and is something we leave to future work. The linear nature of MUSIC makes the model inherently interpretable and can reveal biological pathways driving the observed co-variation patterns via a generalised loadings matrix. To comprehensively reveal non-lineartities in such complex datasets would require to to adapt the factorization process, for example via non-linear kernels. This however, would make interpretation of results challenging.

## Supplementary Information

### 1 Supplementary material

#### 1.1 Full generative model

MUSIC is defined as

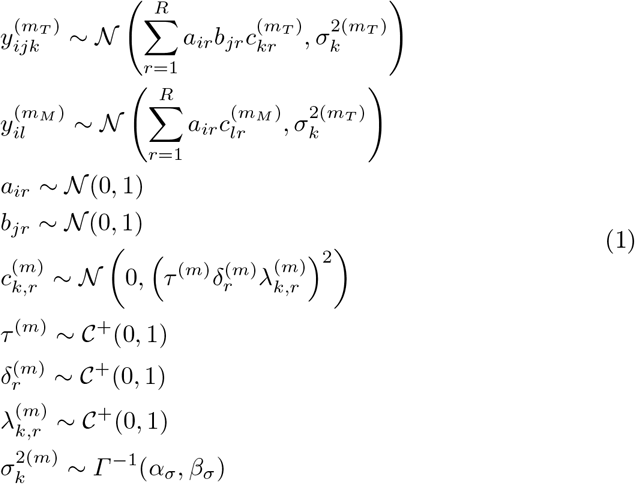

*Γ*^*−*1^(*α, β*) is the inverse-Gamma distribution with shape and scale parameters *α* and *β*. We choose *α* = 1 and *β* = 0.5.

#### 1.2 Joint probability distribution

With *θ* denoting all parameters in the model, the joint probability distribution can be written as:

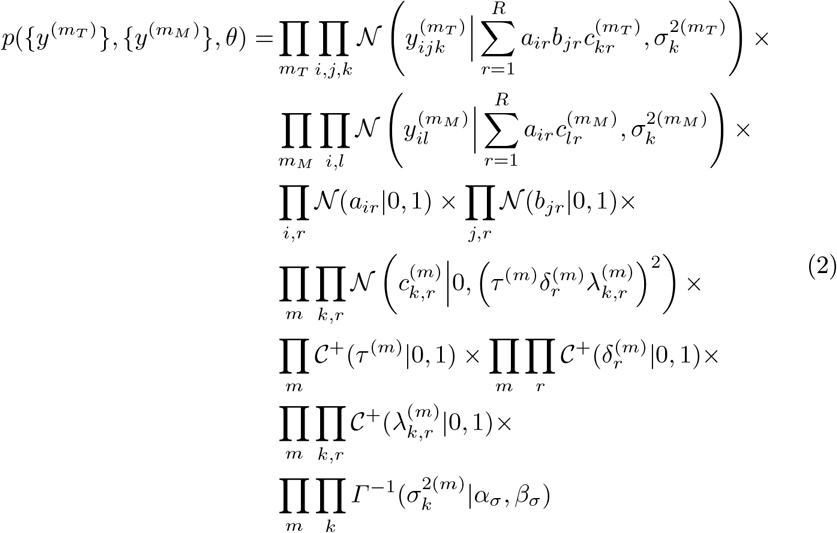

Here, the first line represents the likelihood for all rank-3 tensors, the second line represents the likelihood for all matrices. The third line represents the priors for the shared components **a** and **b**. The fourth line represents the hierarchical prior structure for the view-specific components **c**. The fifth and sixth lines represent the priors for the hierarchical variance component and the last line represents the priors for the observation noise variances.

### 2 Supplementary figures

#### 2.1 Cytokine Data

##### 2.2 Single cell data

**Fig. 1:**
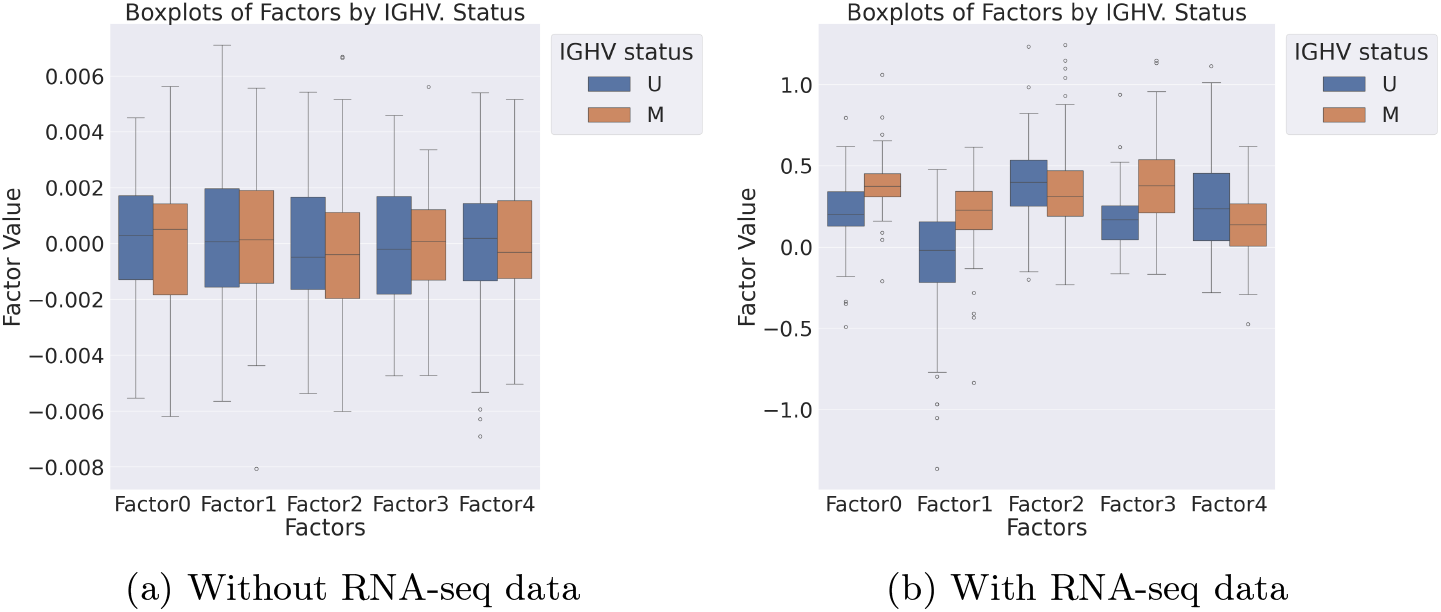
Boxplot of the sample factor values for IGHV status. Stars represented a pvalue < 0.05 at the t-test. The addition

**Fig. 2:**
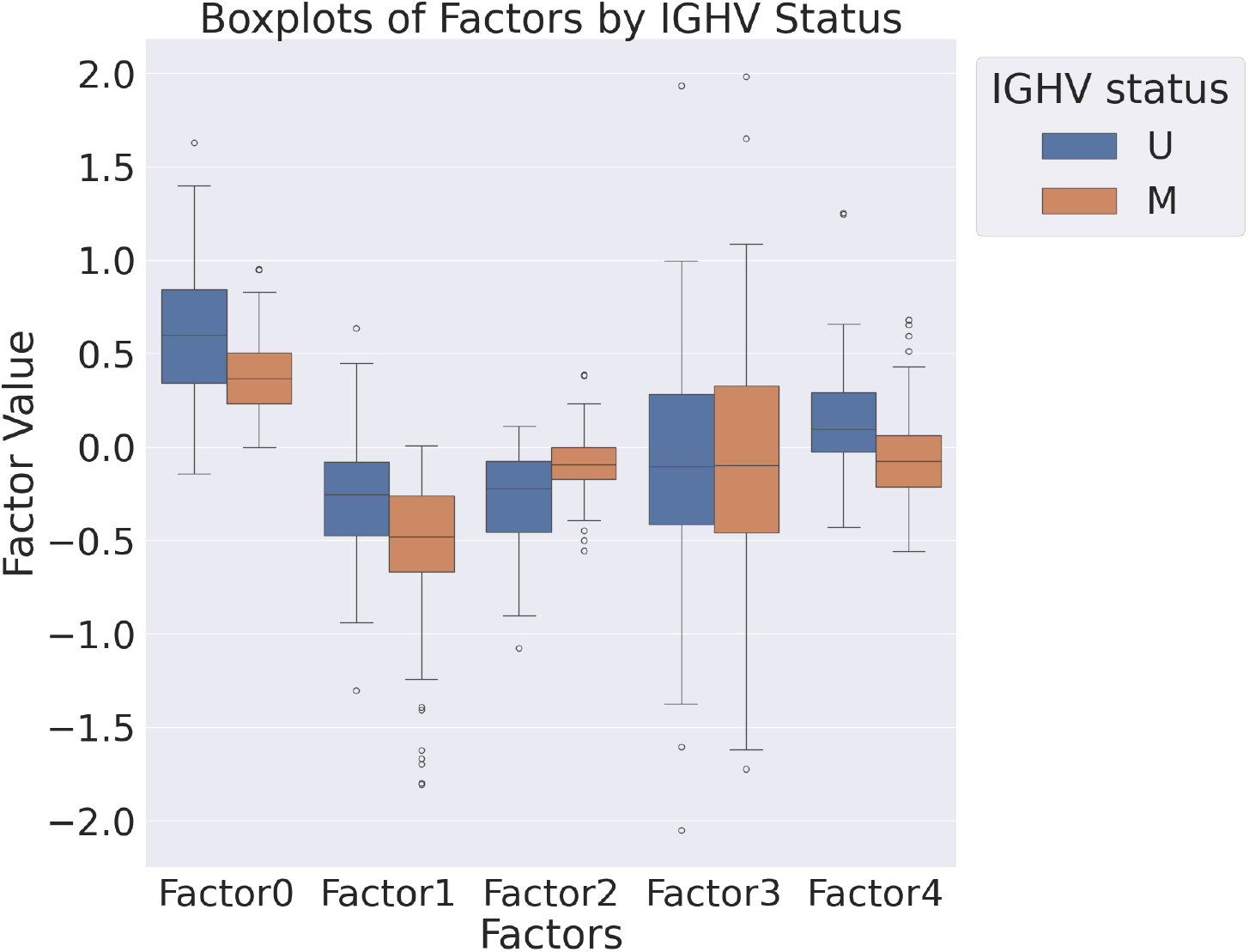
Boxplot of the sample factor values for IGHV status when running Tensorly’s PARAFAC method on the data. Stars represented a pvalue < 0.05 at the t-test

**Fig. 3:**
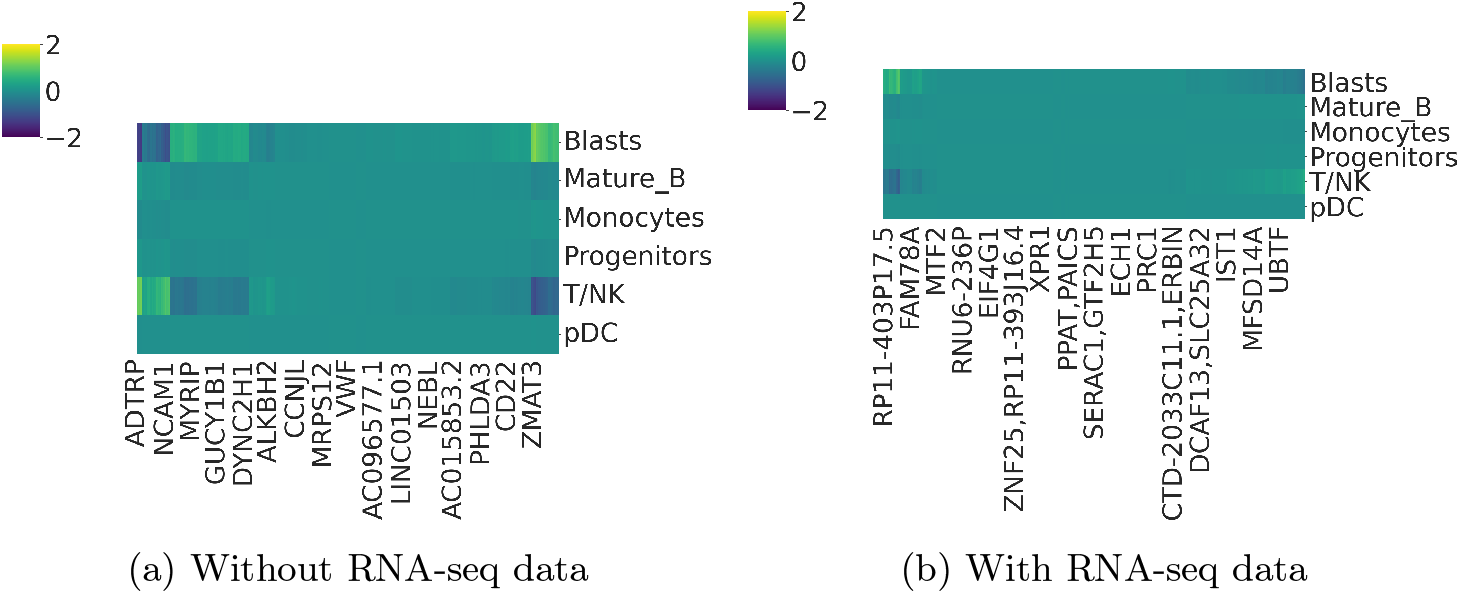
Heatmap of the factor loadings for the patients relative to the drugs for factor 0. The x axis represent the patients

**Fig. 4:**
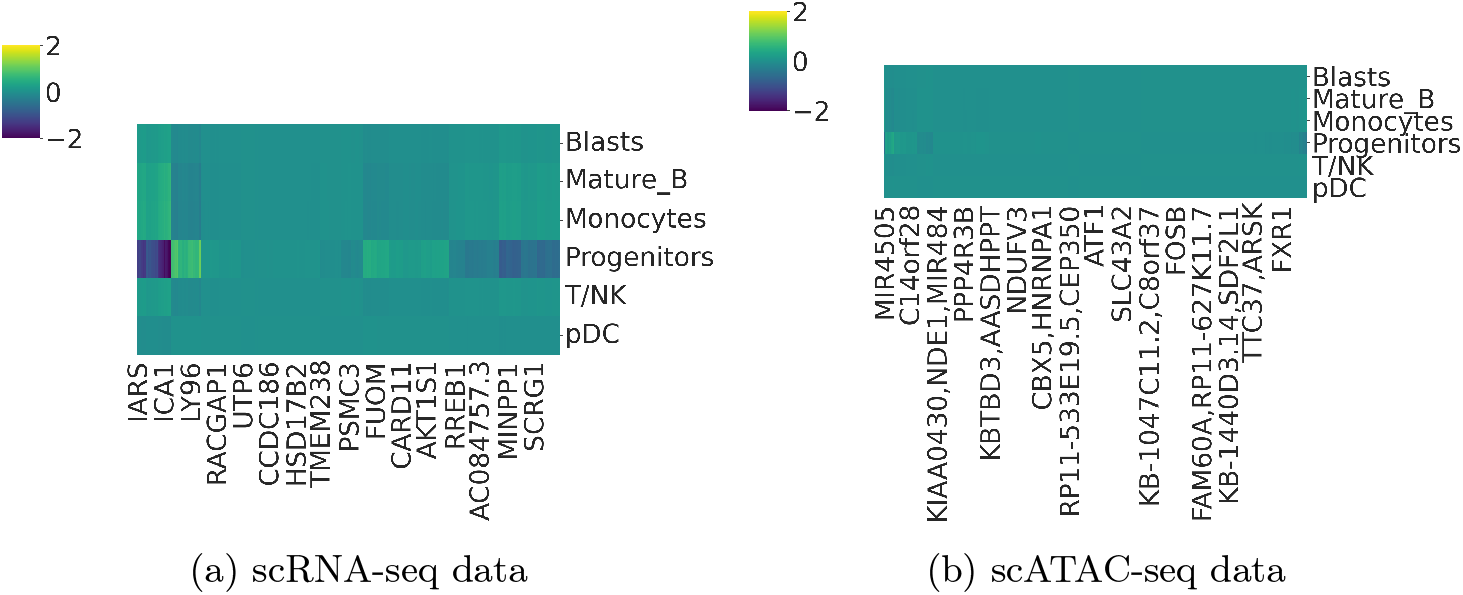
Heatmap of the factor loadings for the cell types relative to the features for factor 3. The x axis represent features.

**Fig. 5:**
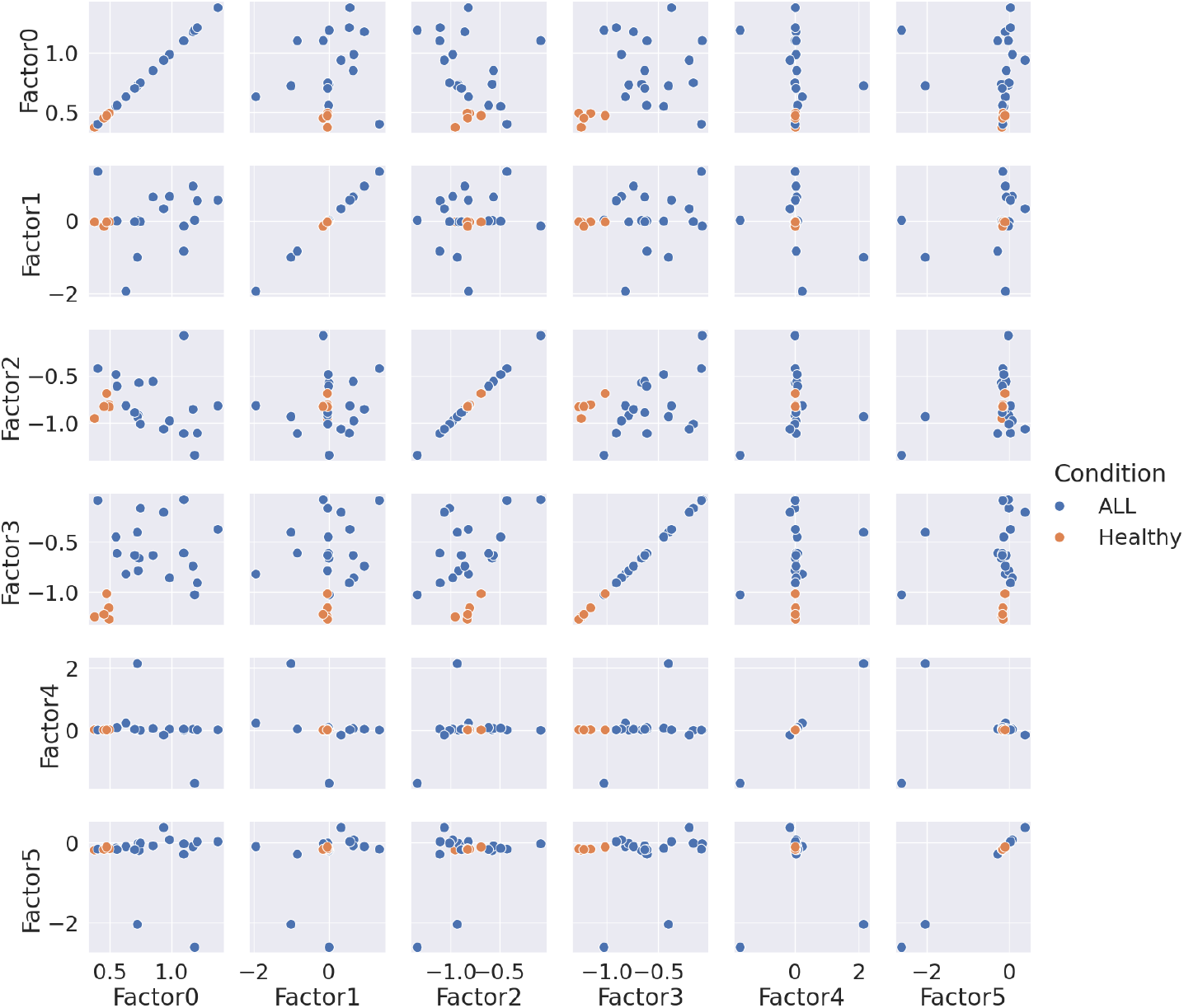
Scatterplot of patients embeddings

## Footnotes

5 We use standard notation where tensors and matrices are denoted with bold capital letters, vectors with bold non-capital letters and scalars with italic non-capital letters.

## References

1. Argelaguet, R., Velten, B., Arnol, D., Dietrich, S., Zenz, T., Marioni, J.C., Buettner, F., Huber, W., Stegle, O.: Multi-omics factor analysis—a framework for unsupervised integration of multi-omics data sets. Molecular systems biology 14(6), e8124 (2018)

2. Bingham, E., Chen, J.P., Jankowiak, M., Obermeyer, F., Pradhan, N., Karaletsos, T., Singh, R., Szerlip, P., Horsfall, P., Goodman, N.D.: Pyro: Deep universal probabilistic programming. The Journal of Machine Learning Research 20(1), 973–978 (2019)

3. Bruch, P.M., Giles, H.A., Kolb, C., Herbst, S.A., Becirovic, T., Roider, T., Lu, J., Scheinost, S., Wagner, L., Huellein, J., et al.: Drug-microenvironment perturbations reveal resistance mechanisms and prognostic subgroups in cll. Molecular Systems Biology 18(8), e10855 (2022)

4. Carvalho, C.M., Polson, N.G., Scott, J.G.: The horseshoe estimator for sparse signals. Biometrika 97(2), 465–480 (2010)

5. Chen, C., Yu, W., Alikarami, F., Qiu, Q., Chen, C.h., Flournoy, J., Gao, P., Uzun, Y., Fang, L., Davenport, J.W., et al.: Single-cell multiomics reveals increased plasticity, resistant populations, and stem-cell–like blasts in kmt2a-rearranged leukemia. Blood, The Journal of the American Society of Hematology 139(14), 2198–2211 (2022)

6. Elizarraras, J.M., Liao, Y., Shi, Z., Zhu, Q., Pico, A.R., Zhang, B.: Webgestalt 2024: faster gene set analysis and new support for metabolomics and multi-omics. Nucleic Acids Research p. gkae456 (2024)

7. Gayoso, A., Steier, Z., Lopez, R., Regier, J., Nazor, K.L., Streets, A., Yosef, N.: Joint probabilistic modeling of single-cell multi-omic data with totalvi. Nature methods 18(3), 272–282 (2021)

8. Ghosh, S., Yao, J., Doshi-Velez, F.: Structured variational learning of bayesian neural networks with horseshoe priors. In: International Conference on Machine Learning. pp. 1744–1753. PMLR (2018)

9. Jayavelu, A.K., Wolf, S., Buettner, F., Alexe, G., Häupl, B., Comoglio, F., Schneider, C., Doebele, C., Fuhrmann, D.C., Wagner, S., et al.: The proteogenomic sub-types of acute myeloid leukemia. Cancer Cell 40(3), 301–317 (2022)

10. Kossaifi, J., Panagakis, Y., Anandkumar, A., Pantic, M.: Tensorly: Tensor learning in python. Journal of Machine Learning Research 20(26), 1–6 (2019)

11. Lee, J., Lyu, H., Yao, W.: Supervised matrix factorization: Local landscape analysis and applications. In: Forty-first International Conference on Machine Learning (2024)

12. Li, Y., Dou, Y., Leprevost, F.D.V., Geffen, Y., Calinawan, A.P., Aguet, F., Akiyama, Y., Anand, S., Birger, C., Cao, S., et al.: Proteogenomic data and resources for pan-cancer analysis. Cancer cell 41(8), 1397–1406 (2023)

13. MacKay, D.J., et al.: Bayesian nonlinear modeling for the prediction competition. ASHRAE transactions 100(2), 1053–1062 (1994)

14. Mani, D., Krug, K., Zhang, B., Satpathy, S., Clauser, K.R., Ding, L., Ellis, M., Gillette, M.A., Carr, S.A.: Cancer proteogenomics: current impact and future prospects. Nature Reviews Cancer 22(5), 298–313 (2022)

15. McKenna, K., Beignon, A.S., Bhardwaj, N.: Plasmacytoid dendritic cells: linking innate and adaptive immunity. Journal of virology 79(1), 17–27 (2005)

16. Mitchel, J., Gordon, M.G., Perez, R.K., Biederstedt, E., Bueno, R., Ye, C.J., Kharchenko, P.V.: Coordinated, multicellular patterns of transcriptional variation that stratify patient cohorts are revealed by tensor decomposition. Nature Biotechnology pp. 1–10 (2024)

17. Qoku, A., Buettner, F.: Encoding domain knowledge in multi-view latent variable models: a bayesian approach with structured sparsity. In: International Conference on Artificial Intelligence and Statistics. pp. 11545–11562. PMLR (2023)

18. Qoku, A., Katsaouni, N., Flinner, N., Buettner, F., Schulz, M.H.: Multimodal analysis methods in predictive biomedicine. Computational and structural biotechnology journal (2023)

